# Distinct susceptibility of masked palm civet nasal epithelium to SARS-CoV-1 and SARS-CoV-2

**DOI:** 10.64898/2026.11.15.766891

**Authors:** Shigeru Fujita, Mizuka Fujiwara, Wilaiporn Saikruang, Ichiro Shimizu, Takenori Ijuin, Masakazu Harunari, Tsutomu Tanikawa, Rima Enomoto, Arnon Plianchaisuk, Shusuke Kawakubo, Takanori Asakura, Kei Sato

**Affiliations:** Division of Systems Virology, Department of Microbiology and Immunology, The Institute of Medical Science, The University of Tokyo, Tokyo, Japan; Department of Biological Sciences, Graduate School of Science, The University of Tokyo, Tokyo, Japan; Yoshida Shodoku Co., Ltd., Tokyo, Japan; Tokyo Pest Control Association, Tokyo, Japan; Ikari Shodoku Co., Ltd., Tokyo, Japan; Division of Pulmonary Medicine, Department of Medicine, Keio University School of Medicine, Tokyo, Japan; Department of Pathology, Immunology and Microbiology, Graduate School of Medicine, The University of Tokyo, Tokyo, Japan; International Research Center for Infectious Diseases, The Institute of Medical Science, The University of Tokyo, Tokyo, Japan; International Vaccine Design Center, The Institute of Medical Science, The University of Tokyo, Tokyo, Japan; Programme in Emerging Infectious Diseases, Duke-NUS Medical School, Singapore

## Abstract

Masked palm civets played an important role in the emergence of SARS-CoV-1, yet physiologically relevant civet respiratory epithelial models remain limited. Here, we establish air-liquid interface cultures from primary masked palm civet nasal epithelial cells. The cultures form a differentiated, barrier-forming epithelium containing ciliated, goblet, and club cell populations and recapitulate key features of native nasal tissue. Using parallel human and civet nasal cultures, we compare infection with SARS-CoV-1 and SARS-CoV-2. In human cultures, both viruses show broadly similar replication kinetics. In civet cultures, SARS-CoV-1 replicates rapidly across donors and causes pronounced epithelial disruption, whereas SARS-CoV-2 replication is slower, less efficient, and more donor dependent. These findings reveal distinct compatibility of the two sarbecoviruses with differentiated civet nasal epithelium and establish a species-matched platform for investigating coronavirus host range and cross-species adaptation.

**Highlights:**

- Civet nasal cultures form differentiated, barrier-forming respiratory epithelium
- SARS-CoV-1 replicates rapidly in civet nasal epithelium
- SARS-CoV-2 replication is slower and varies among civet donors
- SARS-CoV-1 infection disrupts ciliated and mucus-producing epithelial cells

**In Brief:** Fujita et al. establish a differentiated nasal epithelial model from masked palm civets, which were historically linked to SARS-CoV-1 emergence. SARS-CoV-1 replicates rapidly and disrupts the epithelium, whereas SARS-CoV-2 replication is slower and donor dependent, enabling species-matched studies of coronavirus host range and cross-species adaptation.

## Introduction

Cross-species transmission of sarbecoviruses provides a well-documented example of how virus-host compatibility at the respiratory mucosa can shape zoonotic emergence. Severe acute respiratory syndrome coronavirus 1 (SARS-CoV-1; historically SARS-CoV) caused the 2002-2003 SARS outbreak. During the outbreak, SARS-CoV viruses closely related to epidemic human strains were isolated from masked palm civets (*Paguma larvata*) and other mammals in live-animal markets in southern China, and serological studies found evidence of infection among people occupationally exposed to wildlife, particularly civets^1-3^. Subsequent detection of diverse SARS-related coronaviruses in horseshoe bats supported bats as the long-term reservoir, whereas civets are thought to have served as an intermediate or amplification host in market-associated transmission to humans^4-6^.

Consistent with this epidemiological link, masked palm civets are experimentally susceptible to infection with human SARS-CoV-1 isolates and can develop respiratory and systemic pathological changes^7,8^. At the molecular level, angiotensin-converting enzyme 2 (ACE2) is a functional receptor for SARS-CoV-1^9^, and differences in spike-ACE2 compatibility between humans and civets were shown to constitute an important species barrier. Adaptive substitutions in the spike receptor-binding domain can alter the efficiency with which SARS-CoV-1 uses human versus civet ACE2, providing a molecular framework for understanding host adaptation during zoonotic transmission^10-12^. These studies established the importance of civet-virus interactions in SARS-CoV-1 emergence, but much of the mechanistic evidence has come from receptor-binding analyses, engineered cell systems, or whole-animal infection rather than from differentiated primary civet respiratory epithelium.

SARS-CoV-2 is a distinct sarbecovirus, yet it also uses ACE2 for entry and can exploit host proteases such as TMPRSS2 for spike activation^13^. Comparative analyses of ACE2 orthologs have demonstrated that species-specific ACE2 sequence differences can strongly influence SARS-CoV-2 entry^14,15^. In masked palm civets, genomic and transcriptomic analyses have identified expression of ACE2 together with entry-associated proteases including TMPRSS2 and TMPRSS4 in multiple organs^16^. However, the presence of a compatible receptor or entry factor does not by itself define the efficiency, kinetics, or cellular consequences of infection in a differentiated airway epithelium. Such properties can also depend on epithelial cell composition, differentiation state, protease availability, and intrinsic antiviral responses.

Primary airway epithelial cells differentiated at an air-liquid interface (ALI) provide an organotypic system that retains many of these features. Human nasal and airway ALI cultures form a polarized, pseudostratified mucociliary epithelium containing ciliated and secretory cell populations and develop measurable epithelial barrier function^17-20^. These models have been widely applied to respiratory coronavirus infection and can reproduce physiologically relevant viral replication, epithelial cell tropism, apical virus release, and infection-associated epithelial changes that are difficult to assess in conventional immortalized cell lines^21-23^. Accordingly, species-matched ALI cultures offer a useful intermediate platform between reductionist receptor assays and *in vivo* infection studies for dissecting determinants of coronavirus host range.

Despite the central historical association between masked palm civets and SARS-CoV-1, to our knowledge, a differentiated primary civet nasal ALI model has not been reported for direct comparison of SARS-CoV-1 and SARS-CoV-2 infection. In this study, we established an ALI culture system from primary masked palm civet nasal epithelial cells and evaluated its differentiation by histological analysis, mucin staining, transepithelial electrical resistance, and respiratory epithelial lineage markers. We then used this system, together with human nasal ALI cultures, to compare the replication kinetics of SARS-CoV-1 and SARS-CoV-2 and to assess virus-associated changes in differentiated epithelial architecture. This approach was designed to determine whether civet nasal epithelium differentially supports infection of these two sarbecoviruses and to provide a physiologically relevant platform for investigating species- and donor-dependent determinants of coronavirus susceptibility.

## Results

### Establishment of a primary masked palm civet nasal air-liquid interface culture

To establish a physiologically relevant *in vitro* model of the masked palm civet nasal epithelium, we collected primary nasal epithelial cells from masked palm civets captured for wildlife damage management (Fig. 1A). Species identification was first performed on the basis of morphological characteristics and was further supported by mitochondrial cytochrome b (Cytb) sequences assembled from RNA-seq data. The obtained Cytb sequences clustered within a single monophyletic clade together with previously reported *P. larvata* sequences, confirming the species identity of the animals used in this study (Fig. S1).

**Figure 1.**
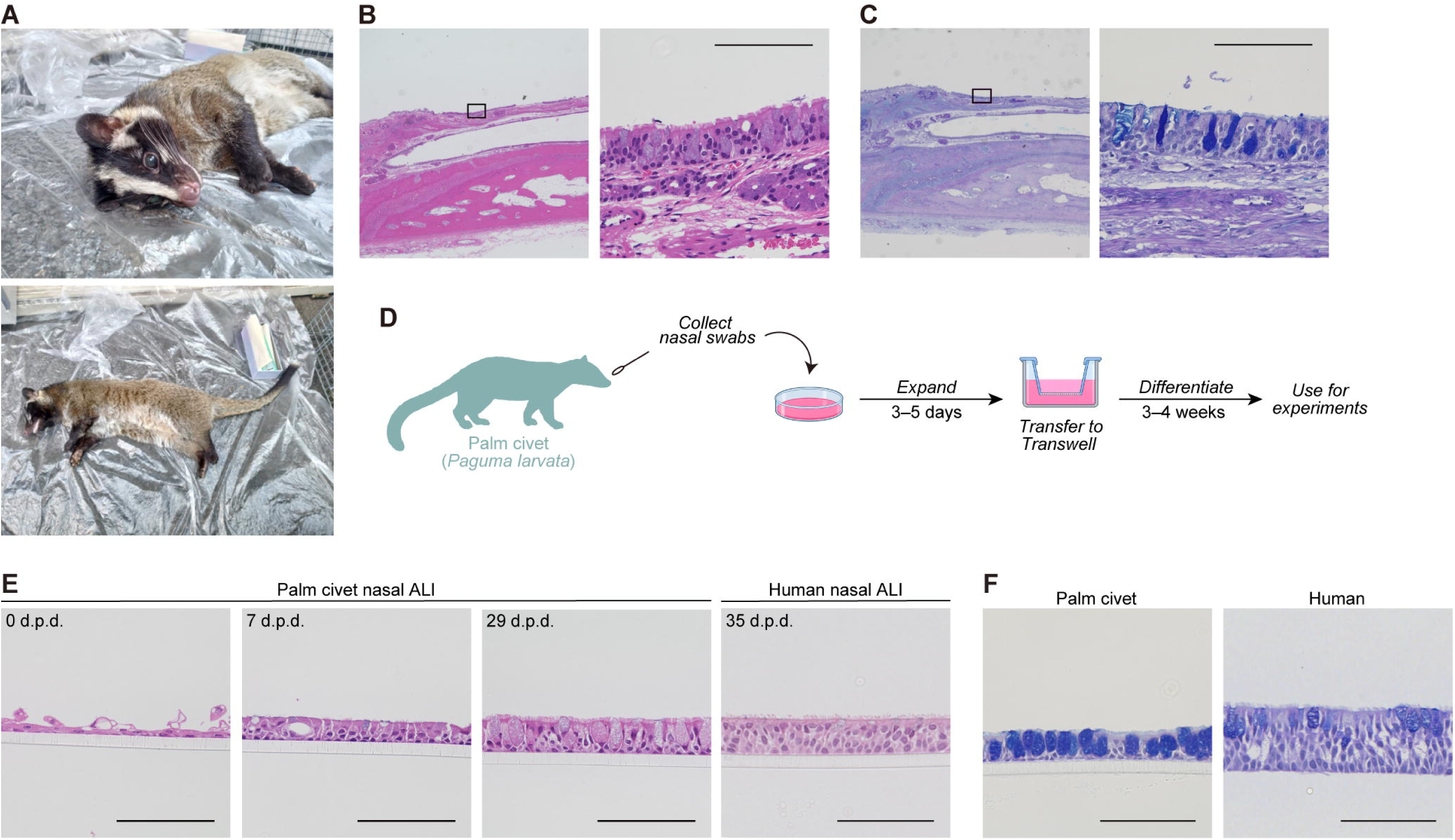
Establishment of masked palm civet nasal epithelial air–liquid interface (ALI) cultures. (**A**) Representative photographs of a masked palm civet (*Paguma larvata*) captured in Japan. (**B** and **C**) Representative hematoxylin and eosin (H&E) staining (**B**) and Alcian blue–periodic acid–Schiff (AB-PAS) staining (**C**) of native masked palm civet nasal tissues. Scale bars, 100 µm. (**D**) Schematic illustration of the establishment of masked palm civet nasal air-liquid interface (ALI) cultures. Nasal epithelial cells were isolated and expanded for 3–5 days, followed by differentiation under ALI conditions for 3–4 weeks. (**E**) Representative H&E staining of masked palm civet nasal ALI cultures at 0, 7, and 29 days post differentiation (d.p.d.). The differentiated human nasal ALI culture also stained at 35 d.p.d. Scale bars, 100 µm. (**F**) Representative AB-PAS staining of masked palm civet nasal ALI cultures at Day 29 (left) and human nasal epithelial ALI cultures at Day 35 (right). Scale bars 100 µm. All data from masked palm civet nasal ALI cultures presented in this figure were obtained from donor PC01. See also **Fig. S1**.

We next examined the histological architecture of native civet nasal tissue. Hematoxylin and eosin (H&E) staining revealed a pseudostratified epithelial structure with prominent cilia on the apical surface, and Alcian Blue-periodic acid-Schiff (AB-PAS) staining visualized mucin-producing goblet cells within the epithelium (Fig. 1B and 1C). These observations established the morphological reference of the native civet nasal epithelium to which the *in vitro* cultures could be compared.

Primary nasal cells obtained by brushing the nasal cavity were expanded in PneumaCult-Ex Plus medium and subsequently seeded onto Transwell inserts for differentiation under ALI conditions (Fig. 1D). At 0 days post-differentiation (d.p.d.), the cells in the ALI culture formed a thin epithelial layer (Fig. 1E). By 7 d.p.d., the epithelial layer had become thicker and sparse cilia were apparent (Fig. 1E). By 29 d.p.d., the culture had developed a substantially thicker, pseudostratified appearance with abundant cilia and goblet-like cells (Fig. 1E). AB-PAS staining at 29 d.p.d. showed abundant mucin-positive cells, confirming the development of a secretory epithelial population (Fig. 1F). Overall, the differentiated ALI cultures recapitulated major morphological features of the native civet nasal epithelium, although goblet-like cells appeared more broadly distributed and rounder in the ALI cultures than in the native tissue (Fig. 1C and 1F), particularly evident in donor PC01, from which the ALI cultures shown in Fig. 1 were generated.

### Functional and molecular validation of differentiated civet nasal ALI cultures

We next assessed epithelial barrier formation and the differentiation of distinct epithelial cell types in ALI cultures generated from independent civet donors. Transepithelial electrical resistance (TEER) increased during differentiation in all three donors examined (PC03, PC06, and PC10). TEER reached a maximum by 7 d.p.d. or 14 d.p.d., depending on the donor, and subsequently remained at relatively stable levels through 28 d.p.d. (Fig. 2A), indicating the formation and maintenance of an epithelial barrier.

**Figure 2.**
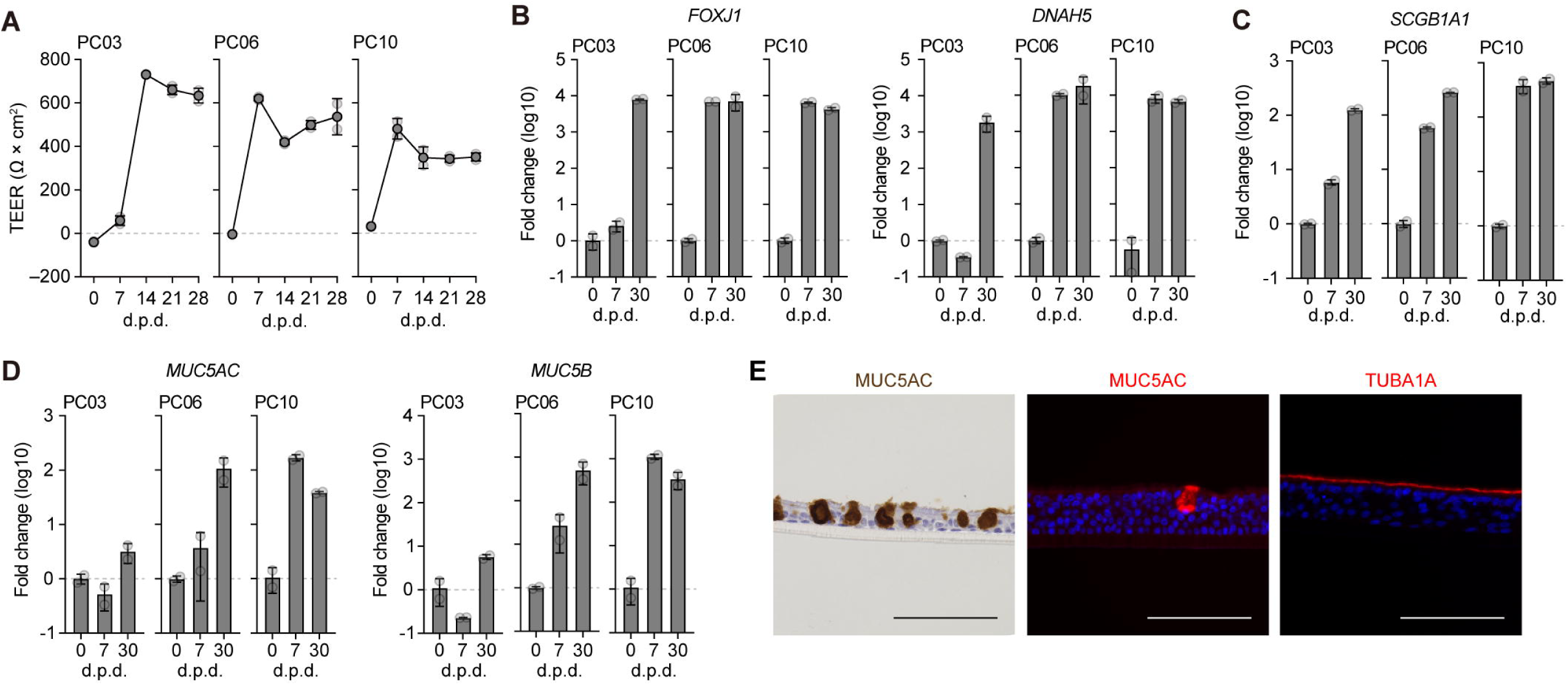
Functional and molecular characterization of masked palm civet nasal epithelial ALI cultures. (**A**) Transepithelial electrical resistance (TEER) was measured at 0, 7, 14, 21, and 28 d.p.d. TEER values are expressed in Ω × cm². Data from three individual masked palm civets, PC03, PC06, and PC10, are shown separately and presented as mean ± SD of two technical replicates for each individual. (**B**–**D**) RT-qPCR analysis of cell-type-specific marker gene expression in masked palm civet nasal epithelial ALI cultures. Ciliated cell markers *FOXJ1* and *DNAH5* (**B**), club cell marker *SCGB1A1* (**C**), and goblet cell markers *MUC5AC* and *MUC5B* (**D**) were analyzed. Expression levels were normalized to those of the 0 d.p.d. sample. Data from three individual masked palm civets are shown separately and presented as mean ± SD of two technical replicates for each individual. (**E**) Representative immunohistochemical staining for MUC5AC and immunofluorescence staining for MUC5AC and acetylated alpha-tubulin (TUBA1A) in masked palm civet nasal ALI cultures. Nuclei were counterstained with DAPI (blue) in the immunofluorescence images. Scale bars, 100 µm. See also **Fig. S2**.

To assess epithelial differentiation at the molecular level, we quantified the expression of respiratory cell markers at 0, 7, and 30 d.p.d. by RT-qPCR (Fig. 2B). Expression of the ciliated-cell markers *FOXJ1* and *DNAH5* increased markedly during ALI differentiation in all three donors, although the timing and magnitude of induction varied among donors (Fig. 2B). The club-cell marker *SCGB1A1* similarly increased after the initiation of ALI culture in each donor and remained elevated at 30 d.p.d. (Fig. 2C). Goblet-cell markers showed greater inter-donor variation (Fig. 2D). *MUC5B* and *MUC5AC* were only modestly induced in PC03, whereas substantially higher expression was observed in PC06 and PC10 during differentiation (Fig. 2D). Thus, differentiation into ciliated and clubcell types was consistently observed across donors, while the extent of goblet-cell differentiation was more donor dependent.

The differentiated epithelial phenotype was further confirmed by immunostaining. MUC5AC-positive goblet cells were readily detected in the differentiated cultures, and acetylated alpha-tubulin (TUBA1A) staining delineated cilia along the apical surface, similar to those observed in human nasal ALI cultures (Figs. 2E and S2). Together with the histological, TEER, and RT-qPCR data, these findings demonstrate that primary civet nasal epithelial cells can be expanded and differentiated under ALI conditions into a structured respiratory epithelium containing major epithelial cell types found in the native nasal mucosa.

### SARS-CoV-1 replicates more rapidly than SARS-CoV-2 in civet nasal ALI cultures

We next investigated whether the established civet nasal ALI culture could be used to compare the replication of SARS-CoV-1 and SARS-CoV-2 in a physiologically relevant respiratory epithelium. Differentiated human and civet nasal ALI cultures were infected with the SARS-CoV-1 Frankfurt 1 strain or the parental SARS-CoV-2 WK-521 strain at an MOI of 0.05, and viral replication was monitored for 7 days (Fig. 3A and 3B). In human nasal ALI cultures, both viruses showed broadly similar replication kinetics, with viral RNA levels and infectious titers increasing after infection and generally peaking around 4-5 days postinfection (d.p.i.), although well-to-well variability was evident in SARS-CoV-2 (Fig. 3A).

**Figure 3.**
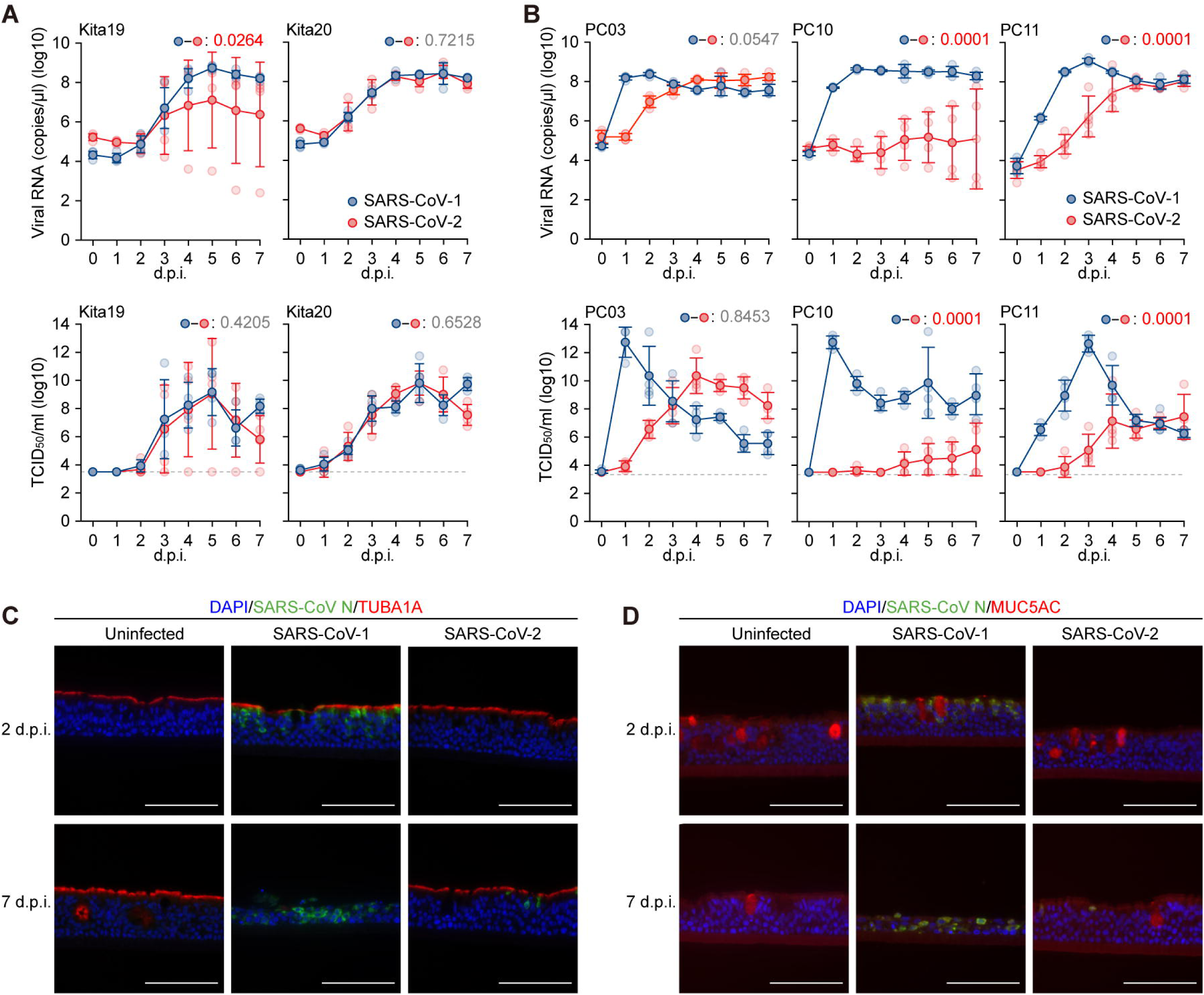
SARS-CoV-1 replicates more efficiently than SARS-CoV-2 in masked palm civet nasal ALI cultures. (**A** and **B**) Viral growth assay. SARS-CoV-1 (Frankfurt 1) and SARS-CoV-2 (WK-521) were inoculated at a multiplicity of infection of 0.05 into human nasal ALI cultures from two individuals, Kita19 and Kita20 (**A**), and masked palm civet nasal ALI cultures from three individuals, PC03, PC10, and PC11 (**B**). The copy numbers of viral RNA in the apical side of cultures were routinely quantified by RT-qPCR (top). The viral titer [50% tissue culture infectious dose (TCID50)] was also measured in the apical side of cultures (bottom). Data from three individual masked palm civets are shown separately and presented as mean ± SD of four technical replicates for each individual. Statistical significance was calculated by permutation test. The familywise error rates (FWERs) calculated using the Holm method are indicated in the figures. (**C** and **D**) Representative immunofluorescence staining of masked palm civet nasal ALI cultures infected with SARS-CoV-2 at 2 and 7 d.p.i. Cultures were stained for SARS-CoV-1/2 nucleocapsid (N) protein (green) and acetylated alpha-tubulin (TUBA1A) (red) (**C**) or MUC5AC (red) (**D**). Nuclei were counterstained with DAPI (blue).

A different pattern was observed in civet nasal ALI cultures. SARS-CoV-1 replicated rapidly in all three civet donors examined (PC03, PC10, and PC11), with viral RNA and infectious titers increasing sharply during the first several days after infection and generally reaching high peaks by 1-3 d.p.i. (Fig. 3B). By contrast, SARS-CoV-2 replication was delayed and more variable among donors. In PC03 and PC11, SARS-CoV-2 replication increased over several days but lagged behind that of SARS-CoV-1. In PC10, SARS-CoV-2 replication was particularly inefficient, and productive replication was not detectable in some wells. Thus, despite donor-dependent variation, civet nasal ALI cultures consistently supported more rapid and robust early replication of SARS-CoV-1 than of SARS-CoV-2.

To visualize infected cells and assess changes in epithelial architecture, we performed co-immunostaining for viral nucleocapsid protein and respiratory epithelial markers in civet ALI cultures (Fig. 3C and 3D). At 2 d.p.i., SARS-CoV-1 nucleocapsid-positive cells were widely detected, whereas SARS-CoV-2 nucleocapsid staining was not detectable. By 7 d.p.i., SARS-CoV-2 nucleocapsid-positive cells became detectable but remained less abundant than those in SARS-CoV-1-infected cultures. SARS-CoV-1 infection was also accompanied by an apparent reduction in acetylated alpha-tubulin and MUC5AC staining and by thinning of the epithelial layer relative to mock-infected and SARS-CoV-2-infected cultures. These observations are consistent with the replication data and indicate that SARS-CoV-1 productively infects civet nasal epithelium earlier and is associated with more pronounced disruption of the differentiated epithelial architecture than SARS-CoV-2 under the conditions tested.

## Discussion

In this study, we established a primary nasal ALI culture system from masked palm civets and showed that the cultures reproduce key structural and functional features of the native nasal epithelium. During ALI differentiation, the cultures developed a pseudostratified epithelial architecture with abundant cilia and mucin-producing cells, acquired stable epithelial barrier function, and upregulated markers of ciliated, goblet, and club cells. Primary human nasal epithelial cells cultured at an ALI are widely used as organotypic models because they recapitulate major features of the differentiated airway mucosa, including epithelial barrier function and relevant expression of respiratory epithelial factors^19,20^. Our findings extend this approach to the masked palm civet and provide a physiologically relevant system in which virus replication can be assessed in a differentiated respiratory epithelium from a species with a historical association with SARS-CoV-1.

The most notable virological finding was the markedly different replication kinetics of SARS-CoV-1 and SARS-CoV-2 in civet nasal ALI cultures. SARS-CoV-1 replicated rapidly in all civet donors examined, generally reaching high levels within the first 1-3 days after infection, whereas SARS-CoV-2 replication was delayed, less efficient, and more variable among donors (Fig. 3B). The efficient replication of SARS-CoV-1, but not SARS-CoV-2, in civet nasal epithelium is consistent with the role of masked palm civets as an intermediate host for SARS-CoV-1. SARS-CoV-1 viruses closely related to epidemic human strain were isolated from masked palm civets in live-animal markets during the 2002–2003 outbreak^1^, and experimental studies subsequently demonstrated that civets are susceptible to infection with human SARS-CoV-1 isolates and can develop systemic and respiratory pathological changes^7,8^). The present results therefore provide a tissue-level correlate of these *in vivo* observations, showing that differentiated civet nasal epithelium itself is highly permissive to productive SARS-CoV-1 infection.

The molecular basis for the rapid replication of SARS-CoV-1 in civet airway epithelium remains to be determined. ACE2 is a major determinant of SARS-CoV-1 entry, and previous studies demonstrated that the interaction between viral spike and ACE2 differs between humans and civets and played an important role in the host adaptation of SARS-CoV-1^10-12^. Experimental reconstruction of zoonotic SARS-CoV-1 further showed that adaptation of spike to human airway epithelium can alter its relative fitness on human versus civet ACE2 backgrounds^24^. Importantly, however, the present study used a human SARS-CoV-1 isolate, yet this virus still replicated rapidly in civet nasal ALI cultures. Thus, efficient replication in civet epithelium cannot be interpreted solely as a property of a civet-origin viral spike. In addition to receptor compatibility, civet-specific dependency factors, protease availability, epithelial cell composition, or differences in antiviral responses may contribute to the efficient replication phenotype observed here.

SARS-CoV-2 showed a contrasting phenotype, with substantial donor-to-donor and well-to-well variability and, in some cultures, little or no detectable productive replication. This heterogeneity may be informative rather than merely technical. The civet ALI cultures themselves exhibited donor-dependent differences during differentiation, most clearly in the induction of the goblet-cell markers MUC5AC and MUC5B (Fig. 2D). These observations raise the possibility that the abundance or differentiation-dependent expression of one or more host factors influences SARS-CoV-2 permissiveness. ACE2 and TMPRSS2/TMPRSS4 transcripts have been detected in multiple organs of masked palm civets^16^, and comparative studies have established that species-specific ACE2 properties can strongly influence SARS-CoV-2 entry^14^. Nevertheless, receptor expression alone is unlikely to fully explain the phenotype observed in our differentiated cultures. Our data do not distinguish whether the restriction occurs at viral attachment and entry, at a post-entry replication step, or through differential innate antiviral responses. Identifying the factor or combination of factors that separates SARS-CoV-1-permissive from relatively SARS-CoV-2-restrictive civet epithelial cultures will therefore be an important next step.

The immunostaining analysis further revealed a qualitative difference in the consequences of infection. SARS-CoV-1 nucleocapsid-positive cells were already abundant at 2 days post infection, whereas SARS-CoV-2-positive cells were not readily detected until later (Fig. 3C and 3D). SARS-CoV-1 infection was also associated with reduced ciliated- and goblet-cell marker staining and thinning of the epithelial layer. Previous experimental infection studies reported pathological changes in SARS-CoV-1-infected civets, including viral RNA and lesions in the respiratory tract and other tissues^8^. Our ALI system isolates the respiratory epithelial component of this phenotype and suggests that efficient SARS-CoV-1 replication can directly accompany substantial disruption of differentiated civet nasal epithelium. Whether the loss of epithelial markers reflects preferential infection of particular cell types, direct cytopathic effects, secondary damage to neighboring cells, or altered differentiation states remains unresolved.

Several limitations should be considered. First, the number of civet donors was limited, and the pronounced donor-to-donor variation observed for both epithelial differentiation and SARS-CoV-2 replication indicates that additional animals will be required to define the range of normal biological variation. Second, the comparison was performed with one SARS-CoV-1 strain and one early SARS-CoV-2 strain under a single infection condition; therefore, the observed difference should not be generalized to all sarbecoviruses or to all strains within either virus species. Third, the current data establish a phenotype but do not identify its mechanism. Comparative profiling of host-factor expression across differentiation states and donors, together with analyses of virus entry, replication, and epithelial antiviral responses, should help determine which steps account for the distinct behavior of SARS-CoV-1 and SARS-CoV-2.

In summary, we established a differentiated masked palm civet nasal ALI culture that recapitulates key features of the native respiratory epithelium and supports comparative coronavirus infection studies. The pronounced early replication of SARS-CoV-1, contrasted with the slower and more heterogeneous replication of SARS-CoV-2, is consistent with the distinct historical host relationships of these viruses and highlights the importance of studying viral host range in physiologically differentiated, species-matched epithelial systems. This platform should enable further investigation of the host determinants that govern sarbecovirus replication and cross-species adaptation in the civet respiratory tract.

## RESOURCE AVAILABILITY

### Lead contact

Further information and requests for resources and reagents should be directed to and will be fulfilled by the lead contact, Kei Sato.

### Materials availability

All unique reagents generated in this study are listed in the key resources table and available from the lead contact with a completed materials transfer agreement.

### Data and code availability

Code for civet gene annotation and Cytb gene extraction can be found at the following GitHub repository: https://github.com/TheSatoLab/Civet-nasal-ALI

## Acknowledgments

We thank Japan Institute for Health Security (JIHS), National Institute for Infectious Diseases (NIID), Japan, for providing a clinical isolate of SARS-CoV-2 strain WK-521 (GISAID ID: EPI_ISL_408667) and SARS-CoV-1 strain Frankfurt 1 (GenBank: AY291315). We would also like to thank Dr. Kaoru Usui, Ms. Naoko Misawa and Ms. Eiko Ogawa (The University of Tokyo) for assistance with collecting civet nasal specimens. The supercomputing resource was provided by the Human Genome Center at the University of Tokyo. We thank Dr. Kaoru Usui (The University of Tokyo) for reviewing the manuscript.

This study was supported in part by AMED ASPIRE Program (26jf0126002, to Kei Sato); AMED SCARDA Japan Initiative for World-leading Vaccine Research and Development Centers "UTOPIA" (JP223fa627001, to Kei Sato); AMED SCARDA Program on R&D of new generation vaccine including new modality application (263fa727002, to Kei Sato); AMED Research Program on Emerging and Re-emerging Infectious Diseases (26fk0108690 to Kei Sato); AMED Japan Program for Infectious Diseases Research and Infrastructure (Collaborative Research via Overseas Research Centers) (26wm0225041, to Kei Sato);JSPS KAKENHI Fund for the Promotion of Joint International Research (International Leading Research) (JP23K20041, to Kei Sato);. JSPS Research Fellow DC2 (24KJ0628, to Shigeru Fujita); JSPS KAKENHI Grant-in-Aid for Scientific Research A (JP24H00607, to Kei Sato); JSPS Bilateral Program JPJSBP123456789 and JSPS-VAST Joint Research Program (120259601, to Kei Sato); AIST KAKUSEI project (FY2024-FY2026 to Shusuke Kawakubo); The Grant for International Joint Research Project of the Institute of Medical Science, the University of Tokyo (25-3151 to Takanori Asakura); and The Cooperative Research Program (Joint Usage/Research Center program) of Institute for Life and Medical Sciences, Kyoto University (to K. Sato).

## Author Contributions

S.F. and M.F. collected the specimen from civets.

I.S., T.I., M.H., and T.T. supported the sample collection from civets.

S.F. and M.F. performed civet ALI preparation.

S.F. and W.S. performed infection assay.

S.F., M.F., and W.S. performed immunostaining.

A.P. performed civet gene annotation.

S.K. performed phylogenetic analysis.

S.F. and S.K. performed Cytochrome b gene extraction of civet samples.

T.A. prepared human ALI and supervised civet ALI preparation.

K.S. supervised the project.

S.F. and K.S. wrote original manuscript.

All authors reviewed and proofread the manuscript.

## Methods

### Ethics statement

The protocol involving nasal specimens from human subjects recruited at Kitasato Institute Hospital were reviewed and approved by the Institutional Review Boards of Kitasato Institute Hospital (approval ID: 22025).

Nasal cells of masked palm civets used in this study were collected from individuals captured for wildlife damage management under a permit from the prefectural governor pursuant to The Act on the Protection of Wildlife and Hunting. Consultation with the Institutional Animal Care and Use Committee of the Institute of Medical Science, The University of Tokyo, confirmed that this study does not constitute an animal experiment requiring ethics committee approval, as it utilized specimens obtained post-euthanasia from individuals captured for pest control.

### Biosafety considerations

All experiments using the live viruses in experimental animals were conducted in the advanced BSL-3 facility in of the Institute of Medical Science, The University of Tokyo, Japan. The live viruses were handled exclusively at BSL-3 facilities, with personnel wearing full-body personal protective equipment and N95 masks. All experimental facilities have been approved by the Institute of Medical Science, The University of Tokyo.

### Cell culture

VeroE6/TMPRSS2 cells (VeroE6 cells stably expressing human TMPRSS2; JCRB Cell Bank, JCRB1819)^25^ were maintained in Dulbecco’s modified Eagle medium (DMEM, low glucose) (Wako, Cat# 041-29775) containing 10% fetal bovine serum (FBS, Sigma-Aldrich Cat# 172012-500ML), G418 (1 mg/ml; Nacalai Tesque, Cat# G8168-10ML) and 1% penicillin–streptomycin (PS).

Human nasal epithelial cells (HNECs) and civet nasal epithelial cells were created according to protocol as described below (see “preparation of air–liquid-interface of primary human nasal epithelial cells (HNECs)” and “preparation of air–liquid-interface of primary civet nasal epithelial cells (CNECs)” section)

### Isolation and expansion of primary of primary human nasal epithelial cells (HNECs)

Primary HNECs were collected from healthy human donors by brushing from Kitasato Institute Hospital. Briefly, superficial brushing was obtained from the inferior nasal turbinate under direct vision through a disposable nasal speculum (Matsuyoshi, Cat# ME1102). Both nostrils were brushed 5 times without anesthesia using a sterile plastic brush, Cytopick (Matsunami, Cat# JJ00000). Nasal cells were isolated by digestion with 5 ml of freshly prepared collagenase IV (1 mg/ml) (Thermo Fisher Scientific, Cat# 17104019), DNase (0.1 mg/ml) (Worthington Biochemical Corporation, Cat# LS002139), and 5 ml of Accutase (Nacalai Tesque, Cat# 12679-54) for 10 minutes at room temperature. After neutralization of these enzymes by fetal bovine serum, HNECs were harvested by centrifugation. HNECs were expanded in PneumaCult-Ex Plus medium (StemCell Technologies, Cat# 05040) following manufacturer instructions.

### Isolation and expansion of primary of primary civet nasal epithelial cells

Nasal cells of masked palm civets used in this study were collected from individuals captured for wildlife damage management under a permit from the prefectural governor pursuant to The Act on the Protection of Wildlife and Hunting. The samples were provided by each contractor under the following permits: Permit No. 6-Kan-Ji-Kei-Ho 366 License No. 1438; Provided by Mr. Takenori Ijuin, Yoshida Shodoku Co., Ltd. (January 16, 2025). Permit No. 5-Kan-Ji-Kei-Ho 350; License No. 2004; Provided by Mr. Masakazu Harunari, Ikari Shodoku Co., Ltd. (March 5, 2024).Permit No. 6-Kan-Ji-Kei-Ho 314; License No. 1194; Provided by Mr. Masakazu Harunari, Ikari Shodoku Co., Ltd. (September 4, 2025).

Superficial brushing was obtained from the nasal turbinate from captured and euthanized masked palm civets was scraped using a sterile plastic brush, Cytopick (Matsunami, Cat# JJ00000). The brush was suspended in a 5 mL microtube (BIO-BIK, Cat# ST-500) containing DMEM (high glucose) (Sigma-Aldrich, Cat# 6429-500ML). The tube was then transported to the laboratory on ice.

The tube containing the harvested nasal cells was centrifuged at 1,500 rpm for 3 min, and the supernatant was discarded. Liver Digest Medium (Thermo Fisher Scientific, Cat# 17703034) was added to the tube, and cells were incubated at 37°C for 10 min. After centrifuging again at 1,500 rpm for 3 min, the supernatant was carefully removed without disturbing the cell pellet. The cell pellet was resuspended in PneumaCult-Ex Plus medium (StemCell Technologies, cat# 05040) and seeded onto cell culture plates pre-coated with PureCol (Advanced Biomatrix, Cat# 5005-100ML), which primarily consists of type I collagen. Cells were cultured for approximately 1 to 2 weeks, and primary nasal basal cell were stocked upon reaching confluence.

### Preparation of air–liquid-interface of primary nasal epithelial cells

To develop the ALI culture, 200,000 cells primary nasal epithelial cells (NECs) were cultured on 12-mm transwell inserts with a 0.4-μm pore size (Corning, Cat# 3460), which was coated by Collagen IV (Sigma-Aldrich, Cat#C7521) and placed in 12-well cell culture plates. All wells were maintained in PneumaCult-Ex Plus medium (StemCell Technologies, Cat# 05040) for the first 3–5 days (until confluence is reached) and then start the ALI culture by removing apical medium and add 1 ml PneumaCult-ALI medium (StemCell Technologies, Cat# 05001) to the basal chamber only. NECs were cultured for 3–4 weeks to allow for cell differentiation, which was visually determined by cilia movement (ciliated cells) and mucus production. Media were replaced every 2 to 3 days (1 ml) during the differentiation phase.

### Fixation and paraffin embedding of *in vivo* tissue and ALI cultures

For histological analyses, including H&E and AB-PAS staining, cultured ALI cells were fixed and embedded in paraffin. Briefly, the culture medium was removed, and 4% paraformaldehyde in phosphate buffered saline (PBS) (Nacalai Tesque, Cat# 09154-85) was added to both the apical and basal sides of the Transwell inserts, followed by incubation for 15 min at room temperature. After fixing, the cells were washed three times with PBS. The membranes were then sandwiched using tissue embedding cassette sponges (Kartell, Cat# 02922-04) inside tissue cassettes (Tissue-Tek Uni-Cassette, Sakura Finetek Japan, Cat# 41560) and stored in 70% ethanol. Civet nasal tissue (nasal cartilage) samples were fixed in 10% formalin neutral buffer solution (Wako, Cat# 068-01663) overnight at room temperature, washed three times with PBS, and stored in 70% ethanol. Paraffin block preparation, sectioning, H&E staining, and AB-PAS staining were performed by Morpho Technology (Hokkaido, Japan).

### Immunostaining

For Immunofluorescence analysis, unstained paraffin sections were prepared by Morpho Technology (Hokkaido, Japan). The paraffin sections were deparaffinized using xylene (Nacalai Tesque, Cat# 36611-45). Antigen retrieval was performed using HistoVT One (Nacalai Tesque, Cat# 06380-76) at 90°C for 20 min. Blocking was carried out using Blocking One Histo (Nacalai Tesque, Cat# 06349-64) for 20 min at room temperature. The slides were then incubated overnight at 4°C with the following primary antibodies: mouse anti-acetylated TUBA1A (Santa Cruz Biotechnology, Cat# sc-23950, 1:500), mouse anti-MUC5AC (Invitrogen, Cat# MA5-12178, 1:500), rabbit anti-SARS-CoV-2 N (Sino Biological, Cat# 40588-T62, 1:500). The slides were subsequently incubated with secondary antibodies for 1 h at room temperature using the following antibodies: goat anti-mouse IgG Alexa Fluor 594 (Invitrogen, Cat# A-11032, 1:500), donkey anti-mouse IgG Alexa Fluor 488 (Invitrogen, Cat# A-21206, 1:500). Mounting and DAPI staining were performed using VECTASHIELD Antifade Mounting Medium with DAPI (Vector Laboratories, Cat# H-1200-10). Fluorescence images were captured using a BZ-X800 All-in-One Fluorescence Microscope (Keyence). Immunohistochemistry for anti-MUC5AC (clone 45M1) was performed by Morpho Technology (Hokkaido, Japan).

### Trans epithelial electrical resistance measurement

TEER was measured using an EVOM2 meter (World Precision Instruments) according to the manufacturer’s protocol. Resistance values (Ω) were measured in triplicate, and TEER (Ω•cm²) was calculated by subtracting the average blank value (an insert cup without cells) from the average sample well value and multiplying by the surface area of the insert membrane.

### RNA extraction and RT-qPCR from ALI cultures

Total RNA was extracted from nasal epithelial cells at Days 0, 7, and 30 after starting ALI culture. The culture medium was removed, and the membranes with adherent cells were peeled off from insert cups using tweezers and harvested into Buffer RLT (Qiagen, Cat# 79216) containing 2-mercaptoethanol. Subsequently, RNA extraction was performed using the QIAamp RNA Blood Mini Kit (Qiagen, Cat# 52304) according to the manufacturer’s protocol. To eliminate genomic DNA contamination, DNase I treatment was carried out using the RNase-Free DNase Set (Qiagen, Cat# 79254). cDNA libraries were synthesized using the RNA to cDNA EcoDry Premix (Random Hexamers) (Takara Bio, Cat# 639546) following the manufacturer’s instructions. RT-qPCR was performed as previously described^26-28^. Reactions were set up using Power SYBR Green PCR Master Mix (Thermo Fisher Scientific, Cat# 4367659) with the primers listed in "Design of RT-qPCR primers targeting respiratory marker genes in masked palm civets" section. The PCR assays were run on a CFX Connect Real-Time PCR Detection System (Bio-Rad).

### Design of RT-qPCR primers targeting respiratory cell marker genes in masked palm civets

To design RT-qPCR primers targeting respiratory cell marker genes in masked palm civets, we first performed orthology-based gene annotation because gene annotations for this species were unavailable in public databases. Coding sequences (CDS) and translated protein sequences of masked palm civet were obtained from the GigaScience database^29^. Protein sequences from human (GCF_000001405.40), mouse (GCF_000001635.27), and four other species closely related to masked palm civet, including cheetah (GCF_027475565.1), domestic cat (GCF_018350175.1), striped hyena (GCF_003009895.1), and meerkat (GCF_006229205.1), were obtained from the RefSeq database^30^. Ortholog groups were inferred from these protein sequences using OrthoFinder2^31^. Civet gene names were then assigned based on the names of orthologous genes within the same ortholog group. The CDSs of civet respiratory cell marker genes were subsequently extracted and used for RT-qPCR primer design. Primer information is summarized in Table S1.

### RNA preparation and sequencing

Total RNA was extracted from civet nasal epithelial cells using the QIAamp RNA Blood Mini Kit (Qiagen, Cat# 52304) according to the manufacturer’s protocol. RNA sequencing was performed by Gene Nex (Dojin Group, Kumamoto, Japan) on a NovaSeq X Plus platform (Illumina) in a 150-bp paired-end run, targeting a data yield of approximately 6 Gb raw data per sample. Strand-specific RNA-seq libraries were constructed following poly(A) selection.

### Assembly of cytochrome b sequences from RNA-seq data and species identification

RNA-seq raw reads were quality-filtered and adapter-trimmed using Trim Galore (v2.3.0) (https://github.com/FelixKrueger/TrimGalore). The cleaned paired-end reads were mapped to the reference sequence (NCBI *Paguma larvata* mitochondrion Reference Sequence: NC_029403.1) using Minimap2 (v2.31)^32^. The resulting alignments were directly sorted and indexed using Samtools (v1.24)^33^. Consensus sequences of the cytochrome b (Cytb) coding region for each sample were generated from the sorted BAM files using samtools consensus and converted into FASTA format for downstream analyses. The resulting Cytb sequences were used as queries for a BLASTN search against the NCBI nt database to retrieve homologous sequences. Hits assigned to the families Viverridae and Felidae were retained, and a multiple sequence alignment was generated using MAFFT v7.525^34^ with the --localpair option. A maximum likelihood phylogenetic tree was inferred using IQ-TREE 2 v2.3.6^35^ under the GTR+F+R5 substitution model, which was selected by ModelFinder^36^ according to the Bayesian information criterion, with 1,000 ultrafast bootstrap replicates. The tree was visualized using PearTree (https://github.com/artic-network/peartree).

### SARS-CoV-1 and SARS-CoV-2 preparation and titration

The working virus stocks of SARS-CoV-1 and SARS-CoV-2 were prepared and titrated as previously described^37,38^. In this study, clinical isolates of a parental SARS-CoV-2 WK-521 strain^25^ (GISAID ID: EPI_ISL_408667) and SARS-CoV-1 Frankfurt 1 strain^39^ (GenBank: AY291315.1). In brief, 20 μl of the seed virus was inoculated into VeroE6/TMPRSS2 cells (5,000,000 cells in a T-75 flask). One hour postinfection (h.p.i.), the culture medium was replaced with DMEM (low glucose) (Wako, Cat# 041-29775) containing 2% FBS and 1% PS. At 3 days postinfection (d.p.i.), the culture medium was harvested and centrifuged, and the supernatants were collected as the working virus stock. The titer of the prepared working virus was measured as the 50% tissue culture infectious dose (TCID50). Briefly, one day before infection, VeroE6/TMPRSS2 cells (10,000 cells) were seeded into a 96-well plate. Serially diluted virus stocks were inoculated into the cells and incubated at 37°C for 4 d. The cells were observed under a microscope to judge the CPE appearance. The value of TCID50/ml was calculated with the Reed–Muench method^40^. For verification of the sequences of SARS-CoV-2 and SARS-CoV-1 working viruses, viral RNA was extracted from the working viruses using a QIAamp viral RNA mini kit (Qiagen, Cat# 52906) and viral genome sequences were analyzed as described below (see “Viral genome sequencing” section).

### Viral genome sequencing

Viral genome sequencing was performed as previously described^41,42^. Briefly, the virus sequences were verified by viral RNA-sequencing analysis. Viral RNA was extracted using a QIAamp viral RNA mini kit (Qiagen, Cat# 52906). The sequencing library employed for total RNA sequencing was prepared using the NEB Next Ultra RNA Library Prep Kit for Illumina (New England Biolabs, Cat# E7530). Paired-end 76-bp sequencing was performed using a MiSeq system (Illumina) with MiSeq reagent kit v3 (Illumina, Cat# MS-102-3001). Sequencing reads were trimmed using fastp v0.21.0^43^ and subsequently mapped to the viral genome sequences of a lineage A isolate of SARS-CoV-2^25^ (strain WK-521, GISAID ID: EPI_ISL_408667) or SARS-CoV-1^39^ (strain, Frankfurt 1, GenBank: AY291315.1) using BWA-MEM v0.7.17^44^. Variant calling, filtering, and annotation were performed using SAMtools v1.9^45^ and snpEff v5.0e^46^. Information on the unexpected substitutions detected is summarized in Table S2).

### Virus infection

In the infection experiments using civet nasal ALI cultures, viral stocks for SARS-CoV-2 (WK-521 strain) and SARS-CoV-1 (Frankfurt 1 strain) were inoculated under the following conditions. Differentiated nasal ALI culture in 12-mm Transwell inserts had approximately 3 million cells/well, and viruses were inoculated onto the apical side at a multiplicity of infection (MOI) of 0.05. The inoculated cells were incubated at 37°C for 1 h. After incubation, the virus inoculum was removed and washed twice with Opti-MEM (Thermo Fisher Scientific, Cat# 11058021). For collection of the viruses, 100 µl Opti-MEM was applied onto the apical side of the culture and incubated at 37°C for 10 minutes, then harvested at the 0 to 7 d.p.i. and used for RT-qPCR to quantify the viral RNA copy number (see “RT-qPCR for viral RNA” section below) and viral titration (see “Viral titration of infection experiments” section below).

### RT-qPCR for viral RNA

RT-qPCR for viral RNA was performed as previously described^37,38^. Briefly, 5 μl culture supernatant was mixed with 5 μl of 2 × RNA lysis buffer [2% Triton X-100 (Nacalai Tesque, Cat# 35501-15), 50 mM KCl, 100 mM Tris-HCl (pH 7.4), 40% glycerol, 0.8 U/μl recombinant Rnase inhibitor (Takara, Cat# 2313B)] and incubated at room temperature for 10 min. Rnase-free water (90 μl) was added, and the diluted sample (2.5 μl) was used as the template for real-time RT-PCR performed according to the manufacturer’s protocol using One Step TB Green PrimeScript PLUS RT-PCR kit (Takara, Cat# RR096A) and the following primers, which were commonly used for SARS-CoV-2 and SARS-CoV-1: Forward N, 5’-CGC ATT GGC ATG GAA G-3’; and Reverse N, 5’-CTC TGT TGG TGG GAA TGT TTT GT-3’. The viral RNA copy number was standardized with a SARS-CoV-2 direct detection RT-qPCR kit (Takara, Cat# RC300A). Fluorescent signals were acquired using a CFX Connect Real-Time PCR Detection System (Bio-Rad).

### Viral titration of infection experiments

The viral titer of the infection experiments was measured as the 50% tissue culture infectious dose (TCID50). Briefly, one day before infection, VeroE6/TMPRSS2 cells (10,000 cells) were seeded into a 96-well plate. Serially diluted each sample (the apical sides of cultures) were inoculated into four lines of the cells and incubated at 37°C for 4 d. The cells were observed under a microscope to judge the CPE appearance. The value of TCID50/ml was calculated with the Reed–Muench method^40^.

### Statistics

In the time-course experiments (Fig. 3A and 3B), we performed a permutation test using the area under the curve (AUC) as a summary measure. We first calculated the observed mean AUC across time points for each condition (i.e., SARS-CoV-1 versus SARS-CoV-2). We then randomized the condition labels at each time point and computed the mean AUC for these permuted datasets. This permutation process was repeated 10,000 times to generate a distribution of mean AUC values under the null hypothesis that the condition has no effect. By comparing the observed mean AUC with the distribution of permuted mean AUC values, we calculated an empirical *P* value for the difference observed between conditions. Subsequently, familywise error rates (FWERs) were calculated by the Holm method. These analyses were performed in R v4.1.2 (https://www.r-project.org/).

## Figure legends

**Figure S1. Maximum likelihood phylogenetic tree of masked palm civet Cytb sequences.**

The tree shows the relationships among PC03, PC10, PC11, and publicly available Cytb sequences from *Paguma larvata* (masked palm civet) and related viverrid species. Felidae was used as the outgroup, and the subtree containing *P. larvata* and closely related species is shown. The *P. larvata* clade is highlighted by a shaded box. Filled tips denote the original samples obtained in this study, and open tips denote sequences retrieved from public databases. Scale axis indicates nucleotide substitutions per site.

**Figure S2. MUC5AC and TUBA1A in human nasal ALI cultures.**

Representative immunohistochemical staining for MUC5AC and immunofluorescence staining for MUC5AC and acetylated alpha-tubulin (TUBA1A) in human nasal ALI cultures. Nuclei were counterstained with DAPI (blue) in the immunofluorescence images. Scale bars, 100 µm.

**Table S1. Summary of Primer information for RT-qPCR analysis of civet nasal ALI, related to Fig. 2.**

**Table S2. Summary of unexpected amino acid mutations detected in the working virus stocks, related to Fig. 3.**

