## Supplementary figures and images for "Distinct susceptibility of masked palm civet nasal epithelium to SARS-CoV-1 and SARS-CoV-2"

### Figure S1

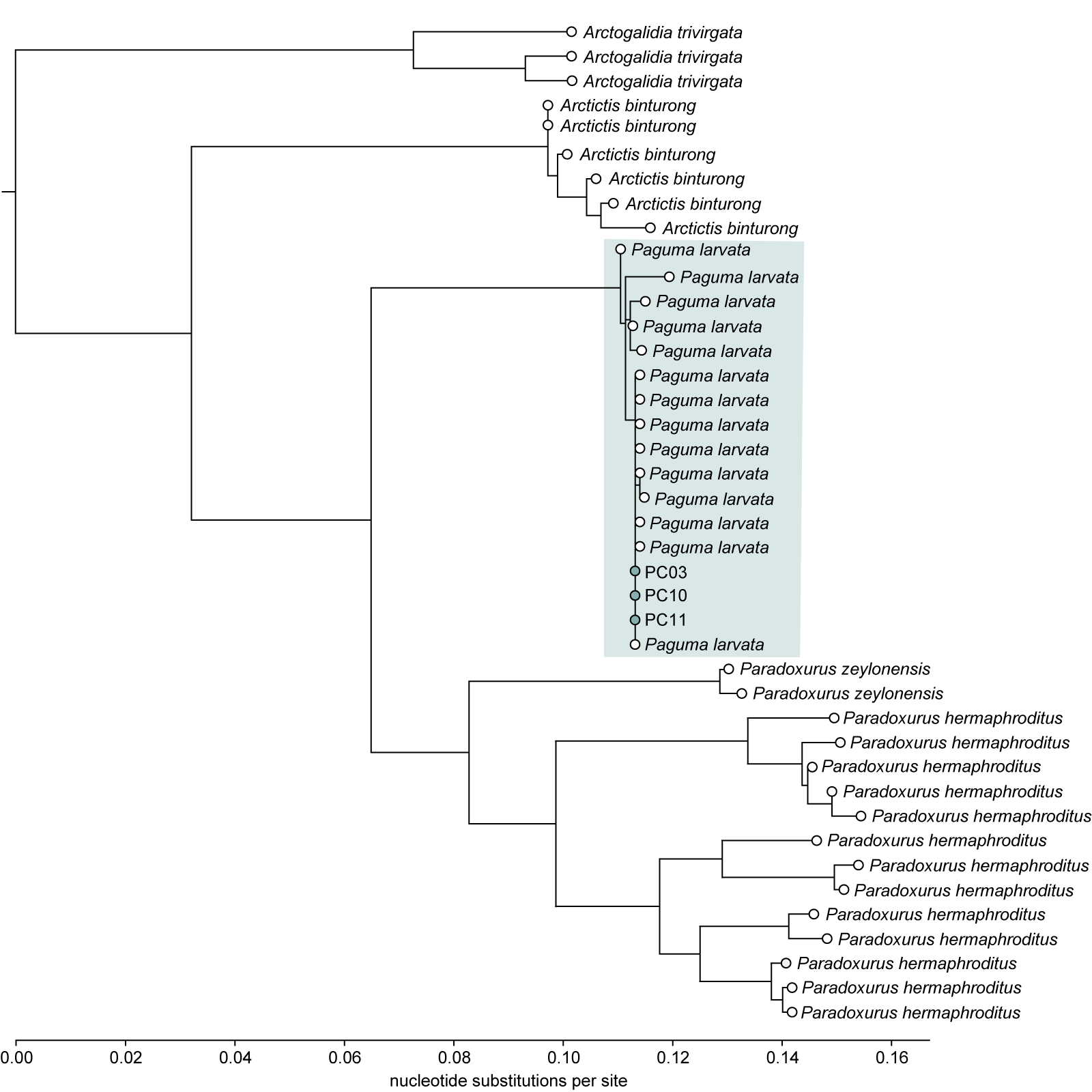

### Figure S2

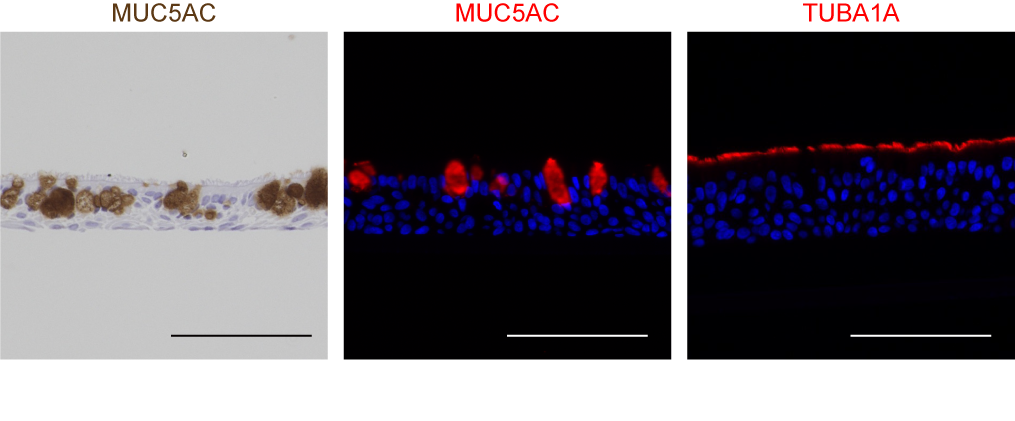
